# Ulcerative colitis is characterized by amplified acute inflammation with delayed resolution

**DOI:** 10.1101/870139

**Authors:** Riccardo Wysoczanski, Alexandra C Kendall, Madhur Motwani, Roser Vega, Farooq Z Rahman, Sara McCartney, Stuart L Bloom, Anna Nicolaou, Derek W Gilroy, Anthony W Segal, Daniel J B Marks

**Affiliations:** Centre for Molecular Medicine, University College London, London, UK; Laboratory for Lipidomics and Lipid Biology, Division of Pharmacy and Optometry, School of Health Sciences, Faculty of Biology, Medicine and Health, University of Manchester, Manchester Academic Health Science Centre, UK; Centre for Clinical Pharmacology and Therapeutics, University College London, London, UK; Department of Gastroenterology, University College London Hospital, London, UK

**Author notes:** Correspondence to Professor Anthony W Segal, Centre for Molecular Medicine, Rayne Building, 5 University Street, London WC1E 6JJ, UK.

## Abstract

The cause of chronic inflammation in ulcerative colitis (UC) is incompletely understood. Here we tested the hypothesis that an excessive acute inflammatory response to bacteria contributes to the pathogenesis. Acute inflammatory responses were provoked *in vivo* in UC patients and healthy controls by intradermal inoculation with bacteria. Vascular responses were quantified by laser Doppler. Inflammatory exudates were recovered in superimposed suction blisters and cells measured by polychromatic flow cytometry, cytokines by multiplex array, and inflammatory lipids by mass spectrometry. Vascular responses in UC patients were heightened at 24h after bacterial injection (p=0·03), and remained abnormally high at 48h (p=0·0005) and this amplified response was seen in UC with Gram-positive as well as Gram-negative organisms (p=0·01). The cellular infiltrate over the injection site, composed largely of neutrophils at 4 hours a was greater in UC (p=0·002). At 48h, the increased numbers of cells in UC were composed of neutrophils (p=0·001) and CD4 lymphocytes (p=0·001). The exaggerated inflammation in UC was not a cytokine-driven phenomenon. Exaggerated onset was normalised in patients taking 5-aminosalicylates, accompanied by increased concentrations of hydroxy fatty acids 9-oxo-octadecadienoic acid (OxoODE; p=0·05) and 13-OxoODE (p=0·01) in resolving exudates. *In vitro*, these compounds suppressed macrophage inflammatory cytokine secretion through PPARγ (p<0·0001). Conversely, 5-aminosalicylates did not inhibit early inflammatory reactions in control participants. Acute inflammatory responses to bacteria in UC are both overly exuberant and slow to resolve. Neutrophils accumulate in excess and persist, in keeping with the pathological appearances of disease flares. These studies also provide new insight into the mechanism of 5-aminosalicylate (5ASA) drugs, which act as pro-resolution rather than indiscriminate anti-inflammatory agents by promoting formation of immunomodulatory hydroxy lipids. While production of these lipids is not defective as part of the underlying disease process, this identifies a novel mechanism of drug action harnessing pro-resolution pathways.

**Summary:** Wysoczanski and colleagues demonstrate that the inflammatory response to injected bacteria is exaggerated and prolonged in ulcerative colitis. This disordered inflammation appears to be associated with increased secretion of PGE2. 5-aminosalicylate drugs, which are used to treat this condition, normalize inflammation and PGE2 secretion, and appear to work through PPARγ

## Introduction

Ulcerative colitis (UC) is characterized by chronic inflammation, principally affecting the large bowel, but with potential to involve skin, eyes and joints (Ungaro et al., 2017). The incidence is 6·3-24·3 per 100,000 person-years (Molodecky et al., 2012), with considerable associated morbidity (including a 30% lifetime risk of surgery and predisposition to colorectal cancer. Eaden et al, 2001). The underlying cause of UC remains unclear, and theories focus on perturbations in the immune system, intestinal barrier or gut contents (including microbiota). Genome-wide association studies (GWAS) have identified over 160 susceptibility loci for inflammatory bowel disease (de Souza and Fiocchi, 2016). Candidate genes have roles in immune responses and mucosal barrier integrity, although mechanistic consequences remain broadly speculative (McGovern et al., 2015). Existing therapies for UC principally target TNF-α, T cells (thiopurines, calcineurin inhibitors and integrin blockers) or janus kinase; however, all have pleiotropic downstream effects, and a substantial proportion of patients do not achieve complete remission and mucosal healing (Ungaro et al., 2017; Verstockt et al., 2018).

An additional aetiological component in UC could be that acute inflammation is excessive. Neutrophils are prominent in the colonic mucosa during flares of UC, and we have previously demonstrated that UC patients mount protracted early inflammation following challenge with *E. coli* (Marks et al., 2006a; Marks et al., 2006b). This could reflect the situation in the large bowel, where there are approximately 10^12^ bacteria/gram of colonic contents and Gram-negative species predominate. While considerable effort has gone into characterizing the pathways and molecules that drive the initiation of inflammation, much less is known about its termination and resolution (Gilroy and De Maeyer, 2015). It is now evident that there are active mechanisms that promote resolution, including clearance of the initiating stimulus, scavenging of cytokines, neutrophil apoptosis, and phagocytosis of effete neutrophils by macrophages. If effective, this coordinated sequence of events prevents the inflammation becoming chronic. We therefore performed a unique set of *in vivo* experiments to delineate the nature of aberrant acute inflammatory responses in UC and to identify the underlying causal processes.

## Results

### Blood flow in response to *E. coli*

We examined acute inflammatory responses following inoculation with UV-killed *E. coli* (UVkEc) in 13 healthy controls (HC), 10 UC patients on no treatment (UC), and 11 UC patients receiving 5ASA drugs (UC-5ASA).

In HC, vascular responses were maximal at 24h, and almost completely resolved within 48h (figure 1A,D). In UC patients not receiving 5ASA drugs, blood flow attained higher peaks (p=0·03), and the rate of change between 24h and 48h (p=0·03; figure 1B,D) again demonstrated that resolution was delayed (Krause et al 1978). Results were comparable whether patients still had their colons *in situ* (n=4) or had previously had a colectomy (n=6). In marked contrast, vascular responses were normal in those patients receiving 5-ASA (figure 1C,D).

**Figure 1:**
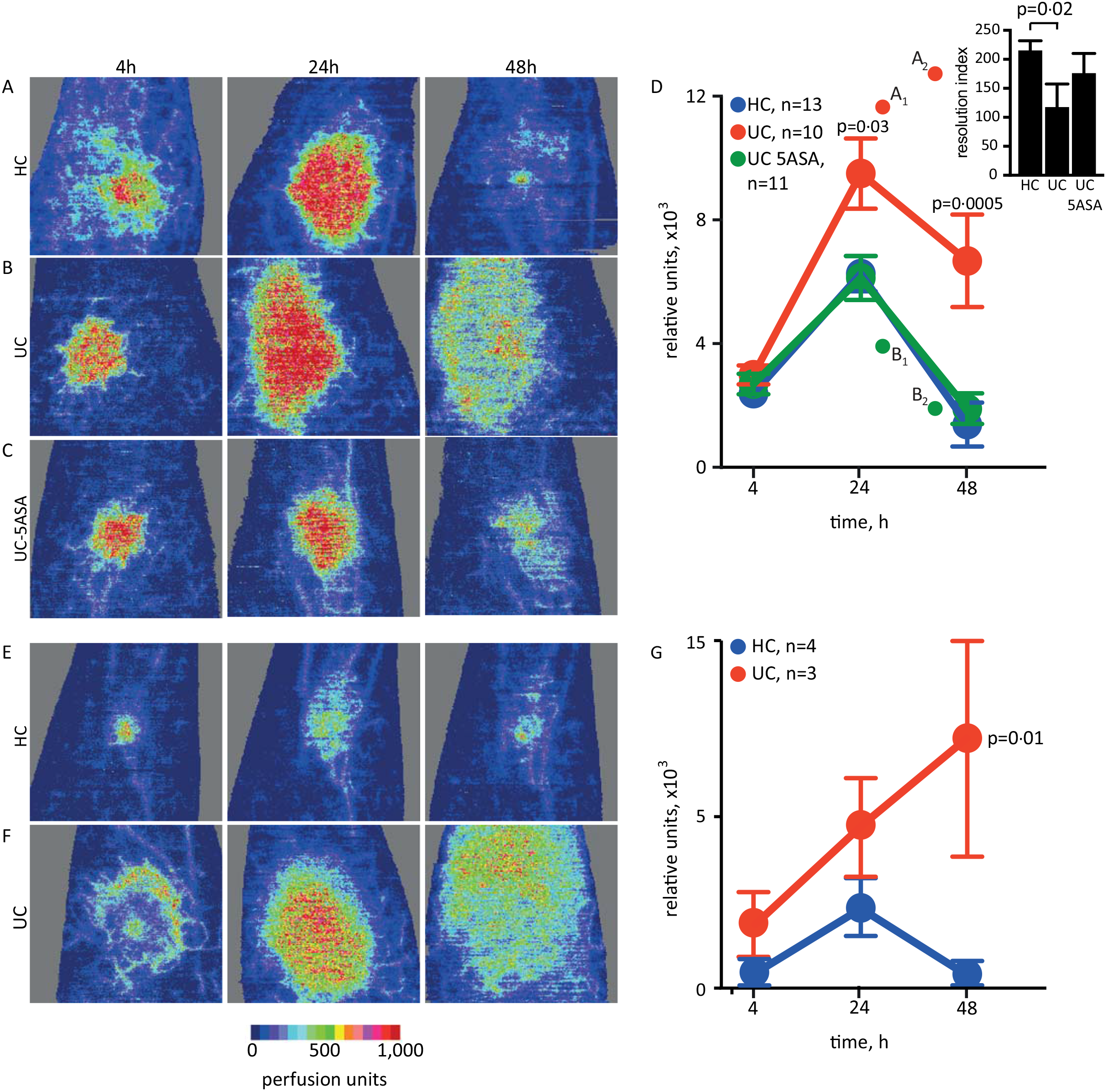
Acute inflammatory responses induced by intradermal inoculation of killed bacteria. (A) Local vascular responses provoked by *E. coli* were observed in HC (n=13) within 4h, peaked by 24h and almost completely resolved within 48h. (B) Responses in UC (n=10) were of greater magnitude, and slower to resolve, except in (C) UC-5ASA (n=11) treatment, in whom blood flow normalised. (D) Blood flow responses to UVkEC were consistently more pronounced and slower to resolve in UC than in HC or UC-5ASA (resolution index, calculated as rate of change of blood flow between 24h and 48h, inset and confirms impaired resolution). One UC patient participated twice, initially off treatment (points A1 and A2) and subsequently while taking 5ASA (B1 and B2). (E) Blood flow responses induced by *S. pneumoniae* followed a similar time course but were less extensive in HC (n=4), but (F) UC (n=3) the enhanced peaks and delayed resolution followed the same pattern. (G) Aggregate responses to UV-killed *S. pneumoniae* confirmed that the abnormality in UC applied to Gram-positive as well as Gram-negative bacteria; at 48h, reactions in UC patients were slowly resolving in two patients but continued to progress in one individual. Data shown as mean±sem.

In addition, during these studies, one UC patient that had been studied when off treatment presented with a flare of their colitis eight months later, and treatment was commenced with 4·8g Mesalazine. We retested this individual 10 months later whilst still on treatment, and found the previously heightened and protracted vascular response had normalized (figure 1D).

### Acute inflammatory reaction triggered by *S. pneumoniae*

To establish whether the enhanced reactions were specific to Gram-negative organisms, we challenged four HC and three UC patients off 5-ASAs with *S. pneumoniae*. The response to this agent was less potent in both HCs and UC patients (figure 1E,G), but the exaggerated early response and delayed resolution were still evident in the UC patients(p=0·02; figure 1F,G).

### Cellular inflammatory response

To identify whether the exaggerated vascular responses were associated with differences in the populations of infiltrating cells, we sampled the acute inflammatory exudates by raising suction blisters over the inoculation sites at 4h or 48h after injection. The former captures initiation of acute inflammation where neutrophil ingress predominates (which is critical for clearance of the bacterial stimulus) (Smith et al., 2009); the latter represents a time at which resolution has occurred in HC but at which inflammation persists in UC. Early inflammatory reactions were hypercellular(p=0·04; figure 2A), due to an approximate doubling of neutrophil numbers (p=0·05; figure 2B); which was normalized in patients on treatment with 5ASA. Early mononuclear phagocyte influx was similar between the groups (figure 2C).

**Figure 2:**
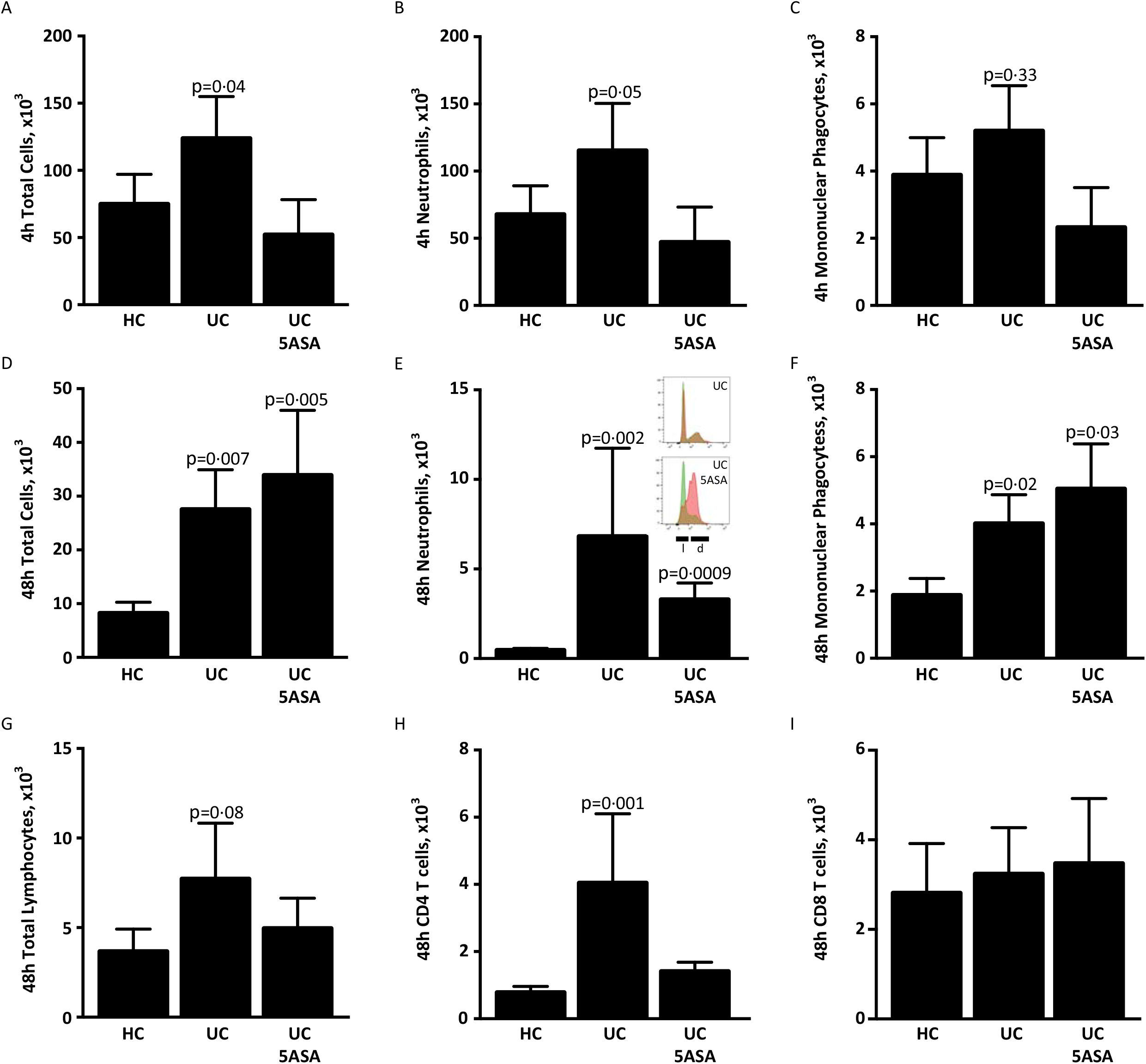
Cellular response to UVkEc. (A) By 4h, UC blisters (n=7) contained approximately double the number of cells compared to HC (n=13); this was normalised in UC-5ASA (n=9). The increased cell numbers were due to expansion of the (B) neutrophil population, whereas (C) the numbers of mononuclear phagocytes were unchanged. (D) In 48h blisters, cells had almost completely cleared in HC, whereas there were excess numbers in both UC groups. (E) There were high numbers of neutrophils in both UC and UC-5ASA participants, but inclusion of a live-dead stain (inset) revealed that in 4h blisters (green) most of these cells in UC (n=3) were alive (l), but by 48h the majority in UC-5ASA (n=2) were non-viable (d). At 48h, (F) mononuclear phagocyte numbers were increased in both UC groups, and (G) there was a trend towards higher lymphocyte numbers in UC patients off treatment; this was due to expansion of (H) CD4 but not (I) CD8 T cells. Data shown as mean±sem.

By 48h, the inflammatory infiltrate had largely cleared in HC (figure 2D), but exudates in UC patients (both off and on treatment) remained hypercellular (p=0·007 and p=0·005, respectively). In patients off treatment there were higher numbers of neutrophils (p=0·002; figure 2E), and mononuclear phagocytes (figure 2F). Although the trend towards expansion of lymphocytes in UC 48h blisters was not statistically significant (p=0·08; figure 2G) there was a clear increase in CD4^+^ T cells (p=0·001; figure 2H), with no differences in CD8^+^ (figure 2I) or B cells. Additional phenotyping identified a CD25^hi^CD127^lo^CCR7^−^CD45RO^+^ dominant subset of the CD4^+^ T cells (supplementary figure 1A,B), which correspond to highly proliferative memory Tregs that preferentially localize to skin (Booth et al., 2010).

In UC-5ASA, at 48h there was an expansion of mononuclear phagocytes compared to HC (p=0·03; figure 2F), many of these cells exhibited a resolution macrophage phenotype (characterized by CD163 scavenger receptor expression; supplementary figure 1C) (Jenner et al., 2014). Neutrophils were also elevated (p=0·0009); however, in three patients off treatment and two UC-5ASA, a viability stain was included and demonstrated that, in UC at 48h, 76·2% of neutrophils were alive compared to only 18·8% in UC-5ASA (figure 2E, inset). Corresponding values in 4h neutrophils were 90·4% and 85·3%, respectively. In addition, 48h neutrophil numbers correlated strongly with mononuclear phagocytes in HC and UC-5ASA (r=0·610, p=0·03 and r=0·952, p=0·001), a characteristic associated with resolving acute inflammation but lost in UC on no therapy (r=0·32, p=0·5).

### Cytokine secretion *in vivo*

Next, we measured concentrations of 29 cytokines in blister exudates. Only interferon-γ was significantly different at 4h, being higher in HC (p=0·04; figure 3A). At 48h, macrophage-derived chemokine (MDC), which contributes to Treg accumulation (Hao et al., 2016), was elevated in HC (p=0·05; figure 3B). After accounting for multiple testing, these findings argue against both the excess production of cytokines, or diminished scavenging of these mediators, as being responsible for the increased inflammation, unless non-classical mediators are involved.

**Figure 3:**
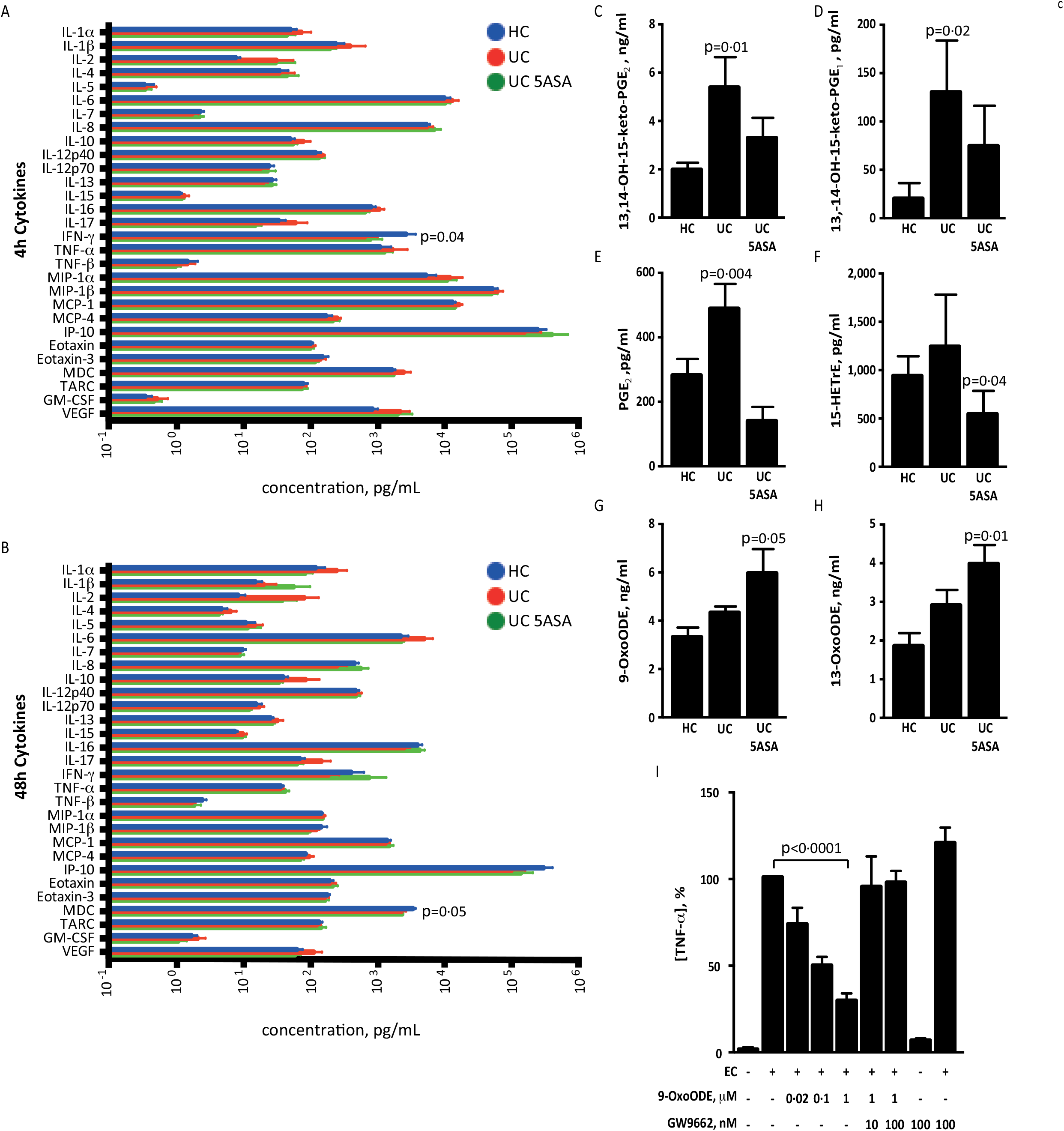
Inflammatory mediators triggered by UVkEc. Cytokine concentrations in (A) 4h and (B) 48h blisters, assayed from 13 HC, 7 UC and 9 UC-5ASA. Lipidomic analysis revealed increased concentrations of (C) PGE_2_ and (D) PGE_1_ metabolites in 4h blisters in UC. (E) In separate experiments, this was recapitulated *in vitro* in peripheral blood monocyte-derived macrophages stimulated with UVkEC from 5 HC, 6 UC and 13 UC-5ASA. In UC-5ASA, (F) HETrE was also reduced in 4h blisters, whereas production of (G) 9-OxoODE and (H) 13-OxoODE was enhanced in 48h blisters. (I) 9-OxoODE caused a dose-dependent suppression of TNF-α secretion from *E. coli* (EC)-stimulated macrophages *in vitro* (n=3 per condition). This was fully reversed by co-incubation with the PPARγ antagonist GW9662. Data shown as mean±sem.

### Lipid mediators *in vivo*

As cytokine profiles did not explain cellular phenotypes, we assayed exudates for inflammatory lipid mediators, which are known to influence both initiation of acute inflammation and its resolution (Stables and Gilroy 2011)Multiple species were detected (see supplementary tables 2 and 3, and supplementary figure 2), of which in 4h blisters only 13,14-dihydro-15-keto-prostaglandin E_2_ (p=0·01; figure 3C) and 13,14-dihydro-15-keto-prostaglandin E_1_ (p=0·02; figure 3D) were significantly different, being elevated in UC and returning towards normal in UC-5ASA. These metabolites are the most stable products of PGE *in vivo* enzymatic degradation, and most reflective of overall production (Samuelsson and Green, 1974).

### Prostaglandin production *in vitro*

Since PGE_2_ can be generated by neutrophils (Weissmann et al, 1982), we considered whether the observed increase might simply reflected elevated cell numbers. Consequently, we assayed PGE_2_ production following exposure to UVkEc in peripheral blood-derived leukocytes *in vitro*. We were unable to detect any generation by neutrophils from any subject, but when we examined macrophages, the pattern of secretion mirrored that observed in the *in vivo* exudate fluid with elevated levels of PGE2 in untreated UC (p=0·004; figure 3E) which reverted to normal in those patients treated with 5ASA.

### Role of OxoODE in resolving inflammation in UC-5ASA

In UC-5ASA, lipid mediators in 4h blisters were comparable to HC, although 15-hydroxyeicosatrienoic acid (HETrE) was marginally lower (p=0·04; figure 3F). In contrast, at 48h, their exudates contained higher concentrations of linoleic acid-derived hydroxy fatty acids 9-oxo-octadecadienoic acid (9-OxoODE; p=0·05; figure 3G) and 13-OxoODE (p=0·01; figure 3H). As the hydroxyoctadecadienoic acid (HODE) precursors to these molecules are elevated during inflammation resolution (Tam et al., 2013), we assessed their effects on cytokine secretion by cultured macrophages, at concentrations comparable to those detected *in vivo*. Co-incubation with UVkEc and increasing concentrations of 9-OxoODE led to dose-dependent suppression of TNF-α secretion (IC_50_ ≈100nM; figure 3I), with no impact on cell viability. Similar results were obtained with 13-OxoODE (supplementary figure 3), although this was less potent. As several oxidized fatty acids covalently bind and activate PPARγ, we examined the impact of GW9662 a potent and selective PPARγ inhibitor(Seargent et al 2004) on TNF-α secretion. This agent completely abrogated the suppressive effects of 9-OxoODE on TNF-α secretion at concentrations of 10nM and 100nM (Figure 3I).

### Differential impact of 5ASA on HC

Given the observed effects of 5ASA on inflammation in UC, we investigated the effect of this drug on the response of HCs that had been loaded with, and maintained on, slow-release oral mesalazine (HC-5ASA) to inoculation with UVkEC. We observed that treatment with this drug did not induce differences in cutaneous blood flow in the 5 HCs (figure 4A), implying that rather than indiscriminately suppressing the general response, the effect of the 5ASAs in UC is more specifically directed to damping down the abnormal exuberant inflammation. Similarly, there were no differences in inflammatory cell infiltrates at 4h (figure 4B), but 48h reactions were hypercellular (p=0·008; figure 4C), with >90% of neutrophils found to be apoptotic. As in UC-5ASA, cytokines were unchanged at 4h (figure 4D), but by 48h (figure 4E) the levels of IL-13 (p=0·03), TNF-α (p=0·03), MCP-1 (p=0·05), MCP-4 (p=0·003) and eotaxin-3 (p=0·003) were minimally diminished. Measurements of the lipids showed that concentrations of 15-HETrE were elevated (p=0·03; figure 4F) and 19,20-dihydroxydocosapentaenoic acid diminished (p=0·03) at 4h; and at 48h PGF2α was reduced (p=0·03; figure 4G).

**Figure 4:**
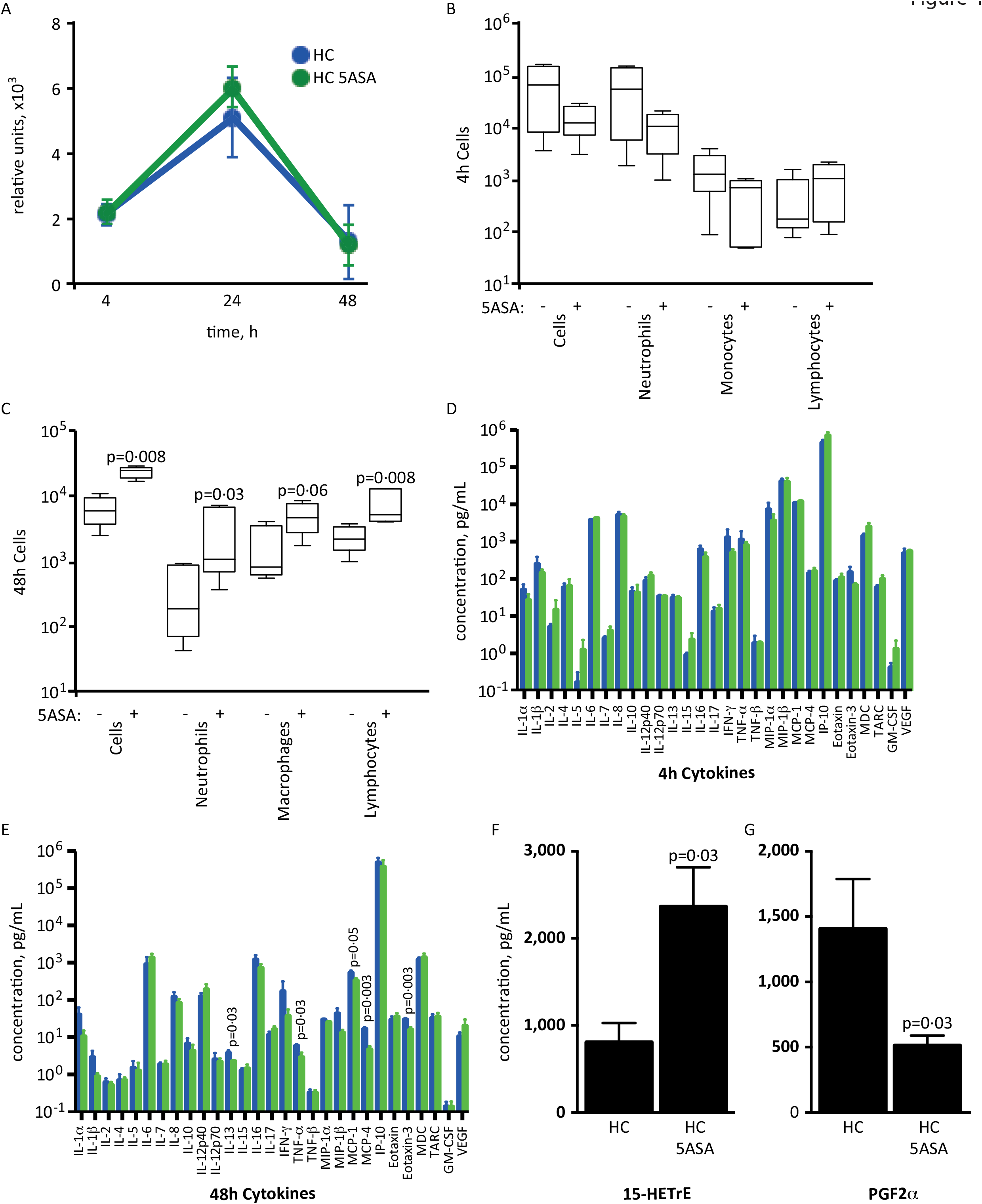
Effects of 5ASA treatment on the acute inflammatory response to UVkEC in HC. (A) Blood flow was unchanged in HC-5ASA subjects (n=5) compared with HC on no treatment (n=5). Blister cellular profiles were no different at (B) 4h, but at (C) 48h all principal cell types were present at greater numbers. Blister cytokine profiles at (D) 4h were unchanged, but (E) subtle variations were seen at 48h. Alterations were also observed in the concentrations of (F) 15-HETrE at 4h, and (G) PGF2α at 48h. Data shown as mean±sem.

## Discussion

These studies demonstrate that the acute inflammatory response is increased in UC, with both an exaggerated onset, and delayed resolution This phenotype contrasts strongly with that observed in Crohn’s disease, where acute inflammation is impaired, which results in delayed bacterial clearance and chronic granulomatous inflammation (Marks et al., 2006a; Smith et al., 2009, Segal AW 2019). This study demonstrates the hazards of conflating these two very different conditions together under the label of “Inflammatory Bowel Disease”, particularly when investigating causal mechanisms with genetic screens such as GWAS(Huang eat al 2017). Excessive neutrophil recruitment and persistence at sites of bacterial inoculation recapitulate the pathological appearances of inflammation in the bowel in UC. The exaggerated reaction to *E. coli* could well help to explain the pathophysiology of UC. While the epithelium normally provides a highly efficient barrier to the bacteria laden faecal contents in health, insults such as infection or trauma allow luminal contents access to tissues of the bowel wall. Trauma could help to explain the observed distribution of UC, which extends proximally from the anus. Progressive dehydration of stool during transit through the colon increases the solidity of the faeces, and with that, the risk of frictional injury to the epithelial barrier. Alterations in genes that are important for maintaining the integrity of the mucus barrier and mucosa could compromise this barrier function and further increase exposure to enteric organisms (McGovern et al., 2015).

The observation of an exaggerated acute inflammatory response in the skin to both gut-derived Gram-negative and to Gram-positive bacteria, raises the question as to why inflammation principally arises in the gastrointestinal tract in UC. This is probably related to the very large numbers of bacteria present in the bowel, readily available to enter the tissues if the mucosa is breeched. It also implies defects in more generic immune mechanisms, rather than specific pathogen-recognition receptors or antigen-directed lymphocyte responses. Similar results were previously reported following intradermal injection of killed *Streptococcus pyogenes*, which produced exaggerated pustular reactions in UC after 48h (Krause et al., 1978). This exaggerated inflammation may additionally provide causal mechanisms for the previously unexplained extraintestinal manifestations in UC (Greuter TI and Vavricka SR, 2019) to conditions affection the joints, lungs and skin such as pyoderma gangrenosum, bronchiectasis (Mateer et al., 2015), Sweet’s syndrome (Ytting et al., 2005), and cutaneous infection with pustular eruptions (Hara et al., 2000).

The literature on PGE_2_ is divergent, with both pro-inflammatory and pro-resolution effects reported, and it may be regarded as a barometer of inflammation. The heightened production we describe here is consistent with previous observations of increased levels in blood and colonic mucosal macrophages and eosinophils in active and relapsing UC (Carty et al., 2000; Raab et al., 1995). The exogenous administration of PGE_2_ causes diarrhoea and pain, and may be a contributing factor to symptoms of the disease. More recently, phospholipase *PLA2G2E*, a primary regulator of arachidonic acid release for PGE_2_ generation which is upregulated by lipopolysaccharide (Murakami et al., 2002), has been consistently identified among GWAS UC susceptibility loci (McGovern et al., 2010; Silverberg et al., 2009).

5ASA drugs were originally developed without knowledge of their molecular target, and the pathways through which 5ASA exerts its influence on the clinical course of UC have been unclear. These drugs are structurally similar to aspirin and also inhibit cyclooxygenase. Our findings suggest that, by also promoting formation of 9-OxoODE and 13-OxoODE in UC, 5ASA exerts additional immunomodulatory effects through PPARγ. This provides a mechanism for previous reports of loss of their ameliorating effects in murine colitis using PPARγ heterozygous knockout animals (Rousseaux et al., 2005); antineoplastic effects in colorectal cancer cells lines associated with PPARγ-related transcriptional activity (Schwab et al., 2008) and abolition of these *in vivo* by GW9662 (Rousseaux et al., 2013); and preliminary reports of efficacy of PPARγ agonists in murine and human colitis (Celinski et al., 2013; Lewis et al., 2008). We postulate that divergent effects in HC-5ASA likely reflect lower precursor substrate release from cell membranes, resulting in decreased shunting towards hydroxy-octadecanoic acid (HODE) (Murakami et al., 2010).

The principal limitation of this study is that bacterial challenge had to be conducted in the skin, rather than in the gastrointestinal tract. This was considered necessary given the restrictive practicalities of injecting bacteria then sampling in the bowel, as well as potential safety implications of provoking excessive neutrophil recruitment in a location where breaching the mucosal barrier could lead to ongoing exposure and propagated responses (particularly given an extreme reaction seen previously in a UC patient) (Marks et al., 2006b). Likewise, as the experiments required the collection of cells trafficking to inflammatory sites, it would be impractical to recover these cells from specific sites in the bowel.

Several of these findings are immediately amenable to clinical translation. The observations that inflammatory lipid mediators play central roles in the pathogenesis of UC and in the therapeutic effects of 5ASA, the major therapeutic tool in UC, are important for the understanding of this disease and for the development of more effective therapy. The acute inflammatory phenotype identifies new biomarkers that should be evaluated for determination of therapeutic efficacy of established and novel medications. Existing PPARγ agonists can be revisited as therapeutics, while more gut-specific molecules are developed alongside stable 9-OxoODE and 13-OxoODE analogues; these would be predicted to enhance inflammation resolution without increasing susceptibility to infection. Likewise, the potential for bacterial challenge as a diagnostic test in patients with indeterminate inflammatory bowel disease might be useful in distinguishing between UC and Crohn’s disease.

## Materials and methods

### Patients with ulcerative colitis and healthy control participants

Patients with endoscopically and histologically proven UC were identified through the gastroenterology clinics at University College London Hospitals (UCLH). Healthy controls (HC) were identified through the Division of Medicine, University College London (UCL), and were defined as individuals with no history of inflammatory bowel disease or other active inflammation disorder. All patients had quiescent disease, defined as a partial Mayo score of 0 and (where available) no active inflammation on recent endoscopy and serum C-reactive protein ≤5mg/L, but previous left-sided or extensive colitis. Each patient who fulfilled the eligibility criteria was approached regarding their interest in participating in research, with a median time from first screening to participation of 35 days. Partial Mayo scores of 0 were re-confirmed on the day of bacterial injection. No participant had received corticosteroid, immunosuppressant or anti-TNF drug therapy within three months, or non-steroidal anti-inflammatory drugs within one week of participation. For UC-5ASA patients, 5ASA was taken orally, and consisted of one of mesalazine 1.2-4.8g daily (n=8), balsalazide 750mg-3g daily (n=2) or sulfasalazine 2g daily (n=1). Where 5ASA was given to HC, this was provided as Mezavant XL 4·8g orally, taken once a day for five days starting 48 hours prior to bacterial injection. Exclusion criteria included age <18 years, HIV infection, pregnancy, breastfeeding, or personal or family history of pyoderma gangrenosum. No participants had active infection while participating in the study. Studies were approved by the National Research Ethics Service Committees London Harrow (project number 13/LO/1541) and Surrey Borders (project number 10/H0806/115). Written informed consent was obtained from all participants.

### Preparation of bacteria for injection into human participants

Antibiotic-sensitive clinical isolates of *E. coli* strain NCTC 10418 (Public Health England, UK) and *S. pneumoniae* serotype 4 (TIGR4 strain; gift from JS Brown) were used for bacterial injection studies. *E. coli* were grown overnight in Luria Broth (Sigma-Aldrich, MO, USA), and *S. pneumoniae* in Colombia blood agar (E&O Laboratories, UK) followed by Todd-Hewitt broth plus 0·5% yeast extract (Sigma), both at 37°C in 5% CO_2_. Bacteria were washed twice in sterile phosphate-buffered saline (PBS; 2,500g, 20min, 4°C), then plated in a sterile petri dish. Counts were determined by optical density for *E. coli* (OD_600_ = 0·365 = 10^8^ organisms/ml) (Smith et al., 2009), and direct counting through serial dilution and plating for *S. pneumoniae*. Bacteria were killed by exposure to a 302nm UV light source (ChemiDoc; Bio-Rad, UK) for 60 min, washed and resuspended in sterile 0·9% saline at a concentration of 1·5×10^8^/ml. Samples were aliquoted into sterile Eppendorf tubes and frozen at −80°C until use. Non-viability was confirmed in multiple cultures by the UCLH Microbiology Department. Either 1·5×10^7^ *E. coli* or 7·5×10^7^ *S. pneumoniae* were injected into the volar aspect of each forearm. Bacterial numbers for inoculation were determined by dose-response studies in HC, including a safety margin given a previous reaction in UC (Marks et al., 2006b). Blood flow at these sites was subsequently quantified using laser Doppler (Moor LDI-HIR, Moor Instruments Ltd, UK), analysed using Moor LDI software (Version 5), and calculated as the product of mean signal and area.

### Suction blisters

To sample exudates, a 10mm suction blister was raised over inoculation sites, 4h or 48h following injection. A blister chamber connected to negative pressure suction (NP-4, Electronic Diversities Ltd, MD, USA) was securely fastened to the forearm. Negative pressure was applied (starting at 2inHg and escalating at 10min intervals to 5-12inHg) until the epidermis separated, forming a blister covering the surface area of the aperture. Pressure was then reduced to baseline at 1inHg every 5min and the chamber removed. Blisters were pierced on their lateral border using a 26·5G needle and exudates collected into tubes containing 50μl 3% sodium citrate (Sigma) in PBS (Gibco, UK). The blister area was cleaned using 0·5% Cetrimide spray and dressed. Collection tubes were weighed to calculate blister volume.

### Analysis of cells in blister fluid

Exudates were centrifuged at 1,000*g* for 5min at 10°C, after which supernatants were aliquoted and immediately frozen at −80°C. Cell pellets were resuspended in 100μl ACK erythrocyte lysis buffer (Lonza, Switzerland) for 1min, then centrifuged at 1,000*g* for 5min at 10°C. Supernatants were discarded and cells resuspended in 100μl PBS with 5% foetal bovine serum (FBS; Gibco, UK) and 0·1% sodium azide (Sigma). Cell were counted with a manual haemocytometer. Leukocyte subpopulations in blisters and peripheral blood samples were characterized by polychromatic flow cytometry, using a previously described gating strategy (Motwani et al., 2016). For isolation of circulating leukocytes, 4ml blood was collected, erythrocytes lysed with ACK buffer, and cells washed with PBS prior to resuspension in PBS/5% FBS. Cells were incubated with an antibody cocktail (supplementary table 1) for 30min at room temperature in the dark. Stained samples were washed in PBS containing 1% FBS and 2mM EDTA (Sigma, UK), then centrifuged at 800rpm for 5min at 4°C. Cells were resuspended in PBS/5%FBS then fixed in an equal volume of 1% paraformaldehyde and stored in the dark at 4°C prior to analysis on a BD LSR Fortessa^TM^ or LSR2 flow cytometer (BD Biosciences, UK). Data were analysed using FlowJo software (FlowJo LLC, OR, USA). Clearance of cells between two time points was defined as (n_1_-n_2_)/t, where n is the number of cells present at a specified time point, and t the interval between time points.

### Analysis of cytokines and lipid mediators in blister fluid

Blister supernatants were assayed for cytokines using the V-PLEX Human Cytokine 30-Plex Kit (Meso Scale Delivery, MD, USA). Lipid mediators were analysed by ultraperformance liquid chromatography coupled to electrospray ionization tandem mass spectrometry (UPLC/ESI-MS/MS), based on protocols published previously.(Kendall et al., 2015; Massey and Nicolaou, 2013) Samples (30-120μl) were defrosted on ice and adjusted to 15% (v/v) methanol:water. Internal standards (20ng each of PGB_2_-*d*4, 12-HETE-*d*8, 8,9-DHET-*d*11 and 8(9)EET-*d*11) were added to each sample and incubated on ice for 30min. pH was adjusted to 3·0 with 1M HCl.

Acidified samples were immediately applied to preconditioned solid-phase cartridges (C18-E; Phenomenex, Macclesfield, UK), washed with various solvents, and lipid mediators eluted with methyl formate (Massey and Nicolaou, 2013). UPLC/ESI-MS/MS analysis was performed on a UPLC pump (Acquity, Waters, Wilmslow, UK) coupled to an electrospray ionization triple quadruple mass spectrometer (Xevo TQ-S, Waters, Wilmslow, UK). Chromatographic separation was performed on a C18 column (Acquity UPLC BEH, 1·7μm, 2·1×50mm; Waters, Wilmslow, UK). Quantitative analysis was based on multiple reaction monitoring-based assays as previously reported (Kendall et al., 2015; Massey and Nicolaou, 2013). Calibration lines were constructed using commercially available standards (Cayman Chemicals, MI, USA).

### Cytokine secretion by neutrophils and macrophages in vitro

Peripheral blood neutrophils were isolated by mixing 20ml venous blood with 10% dextran (MP Biomedicals, OH, USA) to a final concentration of 1%. This was sedimented for 1h then the top layer centrifuged over Lymphoprep (Alere Technologies, Norway) at 2,000rpm for 10min at 20°C. Supernatant was removed and erythrocytes lysed with ddH_2_O. An equal volume of 1·8% saline (Sigma) was added to restore normal tonicity, then cells centrifuged at 1,500rpm for 5min at 20°C. Cells were washed twice in PBS and resuspended in X-Vivo medium (Lonza, Switzerland) at 10^6^ cells/ml before re-plating in tissue culture 96-well plates.

To create macrophage cultures, peripheral blood mononuclear cells were isolated by centrifugation over Lymphoprep at 2,000rpm for 30min at 20°C, then washed three times with phosphate-buffered saline. Cells were resuspended in RPMI-1640 GlutaMAX supplement medium (Gibco, UK) containing 100U/ml penicillin/streptomycin (Gibco, UK) and plated at 10^7^ cells/ml in tissue culture dishes (Nunc, Denmark). After 90min, medium was exchanged with RPMI-1640 containing 10% FBS, 20mM HEPES (Sigma) and 100U/ml penicillin/streptomycin. These were cultured for 5 days at 37°C, 5% CO_2_, with medium supplemented on day 2. On day 5, macrophages were washed three times in PBS, scraped, resuspended at 10^6^/ml in serum-free X-Vivo medium, and re-plated in 96-well tissue culture plates (Falcon, USA). Cells were allowed to adhere overnight prior to stimulation.

Cultured cells were incubated with or without 10^5^ UV-killed *E. coli*. In some experiments, macrophages were co-incubated with 9-OxoODE, 13-OxoODE, GW9662 (Cayman Chemical) or vehicle controls. PGE_2_ and TNF-α secretion were assessed in culture supernatants by ELISA (R&D Systems, UK). Numbers of viable cells in each well were ascertained using Cell Counting Kit-8 (Sigma-Aldrich). All assays utilised protocols from the manufacturers.

### Statistical analysis

Statistical tests were performed using Graphpad Prism version 7·0 (GraphPad Software, CA, USA). Data are presented as mean ± sem unless otherwise stated. Based on observations in previous studies, a sample size of 10 participants per group would be sufficient to detect clinically significant differences in vascular responses with power >95% (α=0·05, β=0·5) (Marks et al., 2006b; Rahman et al., 2010). The Mann-Whitney U test was used for single comparisons, and Kruskal-Wallis ANOVA with Dunn post-tests or two-way ANOVA for multiple comparisons. Where parametric analyses were used, samples were assessed for normality using the D’Agostino and Pearson normality test. Significance values refer to comparison with HC under the same conditions unless otherwise stated. P values <0·05 were considered significant.

During the study, one blister (48h collection in a UC-5ASA patient) ruptured during application of negative pressure, and therefore data on cells and soluble mediators at this time point from this individual were missing; however, all other measurements from this patient, and all samples from other participants, were included in the analyses.

## Contributions

DJBM, AN, DWG and AWS were responsible for the study design. DJBM, RV, FZR, SM and SLB provided clinical care of the patients, and provided clinical phenotyping. DJBM undertook all experiments in conjunction with other authors as follows: RW performed flow cytometry, multiplex cytokine arrays, cell fractionation and culture, cell stimulation and inhibitor studies, and ELISA; ACK performed mass spectrometry for lipid analysis; and MM assisted in development of the blister model. AN planned the lipidomics study, designed the experiments and was responsible for interpretation of lipidomic data. DJBM drafted the initial manuscript, and all authors participated in critical revision. All authors approved the completed article.

## Acknowledgments

This work was funded by the Wellcome Trust, project reference 100098/Z/12/Z. We gratefully acknowledge Prof JS Brown for provision of the streptococcus strains, and M Bonaiti for microbiology input.

## Supplementary Materials

**Supplementary Figure 1:**
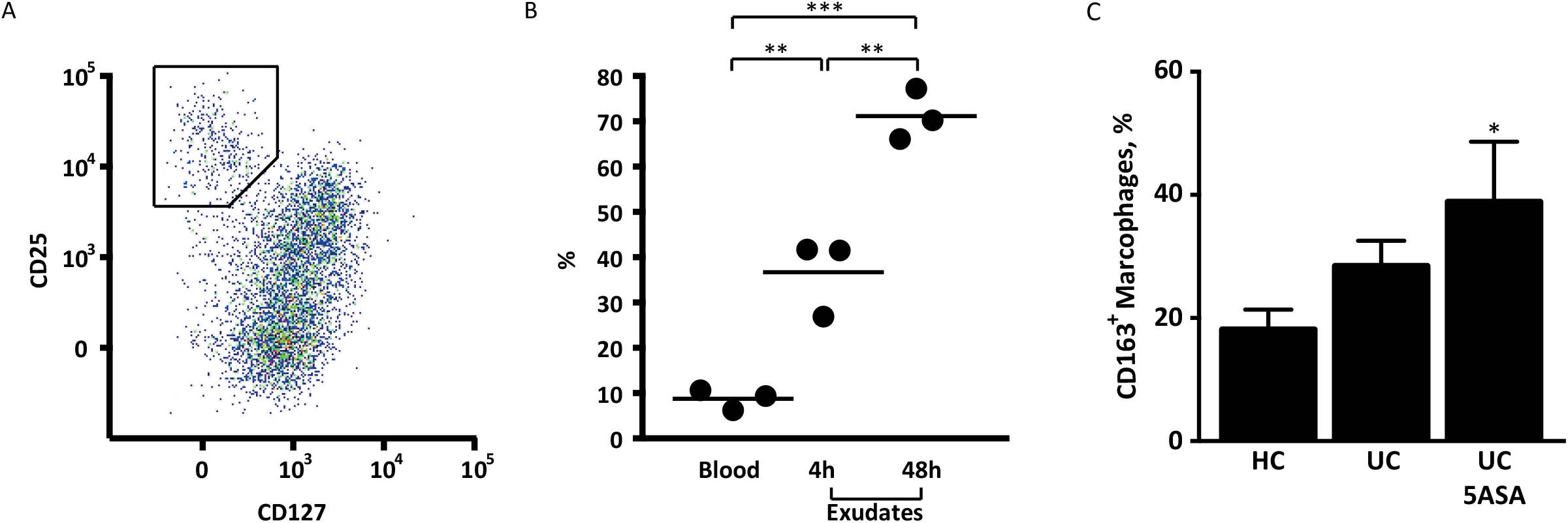
Additional cell phenotyping in blister exudates. (A) Among CD4 lymphocytes in UC patient blood and blisters, a population of CD25^hi^ CD127^lo^ were identified that correspond to a known Treg phenotype; (B) these contributed to a substantial proportion of T cells within UC blisters. (C) In 48h blisters, the proportion of resolution macrophages (as identified by up-regulation of the scavenger receptor CD163) was increased in UC-5ASA. *p<0·05, **p<0·01, ***p<0·001.

**Supplementary Figure 2:**
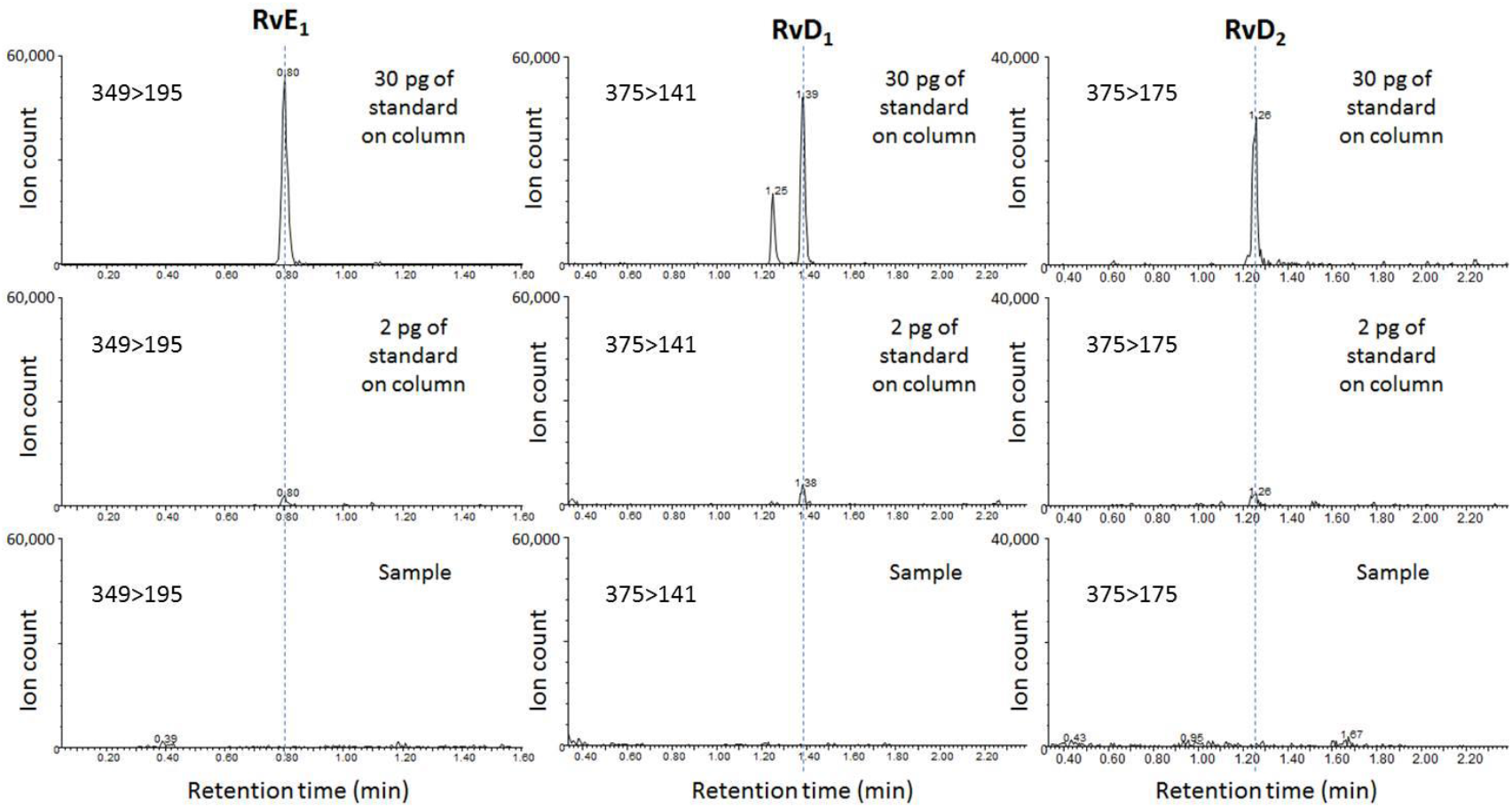
Absence of detectable resolvins in blister fluid. Chromatograms showing an absence of detectable resolvins RvE_1_, RvD_1_ and RvD_2_ in a representative blister fluid sample. Resolvin standards are shown at 30pg on-the-column and 2pg on-the-column, but were undetectable in all samples. The limits of detection for RvE_1_, RvD_1_ and RvD_2_ have been calculated as 180fg/μl, 50fg/μl and 90fg/μl blister fluid, respectively.

**Supplementary Figure 3:**
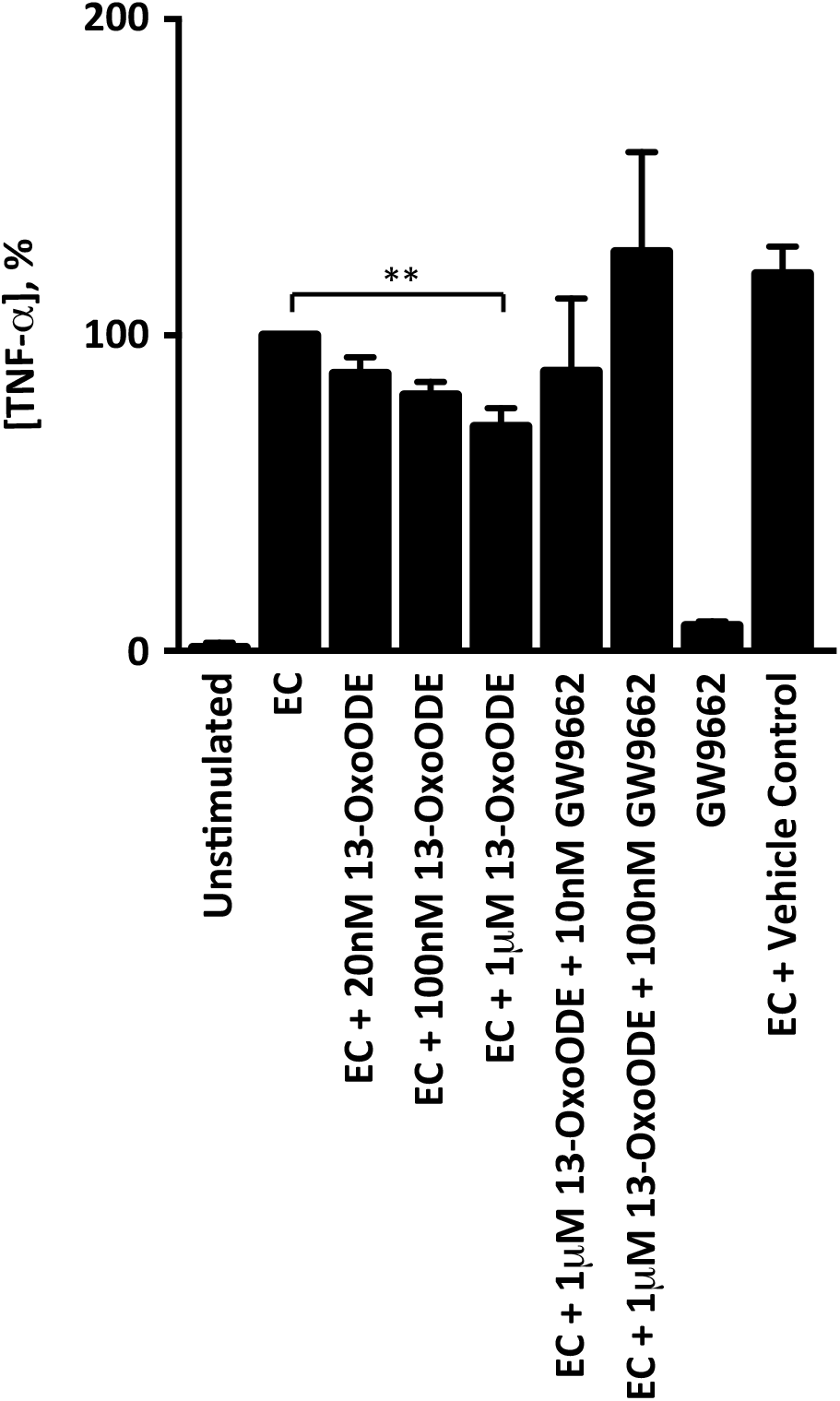
Effect of 13-OxoODE on macrophage TNF-α secretion. 13-OxoODE caused a dose-dependent suppression of TNF-α secretion from *E. coli* (EC)- stimulated macrophages *in vitro*. This was fully reversed by co-incubation with the PPARγ antagonist GW9662. **p=0·01.

**Supplementary Table 1.** Antibody panels used in flow cytometry experiments.

|  | Cell surface marker | Fluorescent conjugate | Antibody clone | Manufacturer |
| --- | --- | --- | --- | --- |
| **Panel 1** |  |  |  |  |
|  | CD3 | FITC | HIT3 | Biolegend |
|  | CD4 | AF700 | RPA-T4 | Biolegend |
|  | CD8 | BV510 | RPA-T8 | Biolegend |
|  | CD14 | BV605 | M5E2 | Biolegend |
|  | CD16 | APC | 3G8 | Biolegend |
|  | CD19 | FITC | HIB19 | Biolegend |
|  | CD45RO | BV785 | UCHL1 | Biolegend |
|  | CD56 | PerCP-Cy5.5 | HCD56 | Biolegend |
|  | CD163 | PE | GHI/61 | BD Biosciences |
|  | CCR7 | BV711 | G043H7 | Biolegend |
|  | HLA-DR | APC-H7 | G46 | BD Biosciences |
|  | CD141 | PE-Cy7 | M80 | Biolegend |
|  | CD1c | BV421 | L161 | Biolegend |
|  | LIVE/DEAD® stain | UV | N/A | Life Technologies |
| **Panel 2** |  |  |  |  |
|  | CD3 | BUV395 | UCHT1 | BD Biosciences |
|  | CD4 | APC | RPA-T4 | BD Biosciences |
|  | CD8 | PerCP-Cy5.5 | RPA-T8 | BD Biosciences |
|  | CD25 | PE | M-A251 | BD Biosciences |
|  | CD45RA | BV605 | HI100 | BD Biosciences |
|  | CD127 | BV421 | HIL-7R-M21 | BD Biosciences |
|  | CD196 | BB515 | 11A9 | BD Biosciences |
|  | LIVE/DEAD® stain | UV | N/A | Life Technologies |

**Supplementary Table 2.** Lipid mediators in 4h blisters. Concentrations in pg/ml (mean ± sem).

|  | HC | UC | UC5ASA | HC 5ASA |
| --- | --- | --- | --- | --- |
| **Eicosanoids** |  |  |  |  |
| PGD2 | 762 ± 210 | 977 ± 233 | 1046 ± 261 | 761 ± 140 |
| PGE1 | 433 ± 66 | 724 ± 199 | 489 ± 176 | 959 ± 408 |
| 13,14-dihydro-15-keto-PGE1 | 20 ± 15 | 130 ± 55 | 74 ± 42 | 52 ± 52 |
| PGE2 | 6573 ± 1282 | 9407 ± 1758 | 11152 ± 4811 | 15336 ± 10053 |
| 13,14-dihydro-15-keto-PGE2 | 1949 ± 271 | 5410 ± 1229 | 3306 ± 816 | 3234 ± 1708 |
| D12 PGJ2 | 1009 ± 111 | 1294 ± 252 | 1192 ± 221 | 1669 ± 371 |
| 15-deoxy-D12,14-PGJ2 | 210 ± 69 | 221 ± 53 | 164 ± 54 | 252 ± 74 |
| PGF1α | 469 ± 68 | 743 ± 226 | 399 ± 106 | 458 ± 52 |
| 6-keto-PGF1α | 2119 ± 372 | 3308 ± 978 | 3781 ± 2059 | 4150 ± 3328 |
| PGF2α | 7335 ± 1110 | 11951 ± 3679 | 7345 ± 2098 | 6093 ± 1986 |
| 13,14-dihydro-15-keto-PGF2α | 0 | 30 ± 20 | 41 ± 28 | 0 |
| TXB2 | 5213 ± 1295 | 7904 ± 1571 | 8957 ± 5593 | 8243 ± 3934 |
| **Hydroxy lipids** |  |  |  |  |
| 4-HDHA | 253 ± 62 | 383 ± 47 | 528 ± 166 | 1137 ± 322 |
| 13-HDHA | 4 ± 4 | 36.55 ± 24.21 | 0 | 0 |
| 14-HDHA | 161 ± 65 | 190 ± 74 | 63 ± 48 | 294 ± 187 |
| 17-HDHA | 0 | 0 | 1055 ± 1055 | 0 |
| 20-HDHA | 18 ± 17 | 6 ± 6 | 21 ± 21 | 0 |
| 11,12-DHET | 160 ± 31 | 243 ± 114 | 143 ± 40 | 169 ± 52 |
| 14,15-DHET | 140 ± 30 | 203 ± 29 | 219 ± 23 | 182 ± 70 |
| 5-HETE | 37 ± 24 | 35 ± 35 | 116 ± 62 | 158 ± 61 |
| 11-HETE | 1047 ± 106 | 1626 ± 576 | 2277 ± 1619 | 1743 ± 1011 |
| 12-HETE | 8237 ± 3071 | 3790 ± 641.5 | 4136 ± 946.1 | 5401 ± 1924 |
| 15-HETE | 2495 ± 442 | 2406 ± 458 | 2829 ± 1876 | 4161 ± 1038 |
| 20-HETE | 410 ± 155 | 584 ± 40 | 922 ± 711 | 735 ± 480 |
| 15-HETrE | 938 ± 202 | 1241 ± 535 | 543 ± 238 | 2370 ± 454 |
| 11(12)-EET | 0 | 72 ± 72 | 154 ± 154 | 46 ± 46 |
| 12-HEPE | 218 ± 164 | 17 ± 17 | 52 ± 33 | 0 |
| 18-HEPE | 234 ± 234 | 0 | 100 ± 63 | 0 |
| 9-HODE | 4744 ± 409 | 6180 ± 1047 | 4621 ± 709 | 5028 ± 600 |
| 13-HODE | 10422 ± 763 | 16192 ± 5145 | 10590 ± 1725 | 13578 ± 1697 |
| 9-OxoODE | 5056 ± 668 | 6382 ± 1461 | 5918 ± 860 | 4987 ± 1208 |
| 13-OxoODE | 3605 ± 733 | 3931 ± 1172 | 3851 ± 685 | 2585 ± 659 |
| 9-HOTrE | 732 ± 135 | 494 ± 115 | 449 ± 181 | 3411 ± 751 |
| 13-HOTrE | 337 ± 54 | 244 ± 87 | 315 ± 118 | 664 ± 139 |
| 9(10)-EpOME | 843 ± 149 | 1371 ± 465 | 1221 ± 311 | 1297 ± 358 |
| 12(13)-EpOME | 1295 ± 219 | 1937 ± 450 | 1883 ± 450 | 2180 ± 459 |
| Trans-EKODE | 1651 ± 243 | 17310 ± 16341 | 1105 ± 139 | 3060 ± 869 |
| 9,10-DiHOME | 1026 ± 84 | 7460 ± 5185 | 1437 ± 326 | 1839 ± 270 |
| 12,13-DiHOME | 1358 ± 254 | 11170 ± 9035 | 2047 ± 495.9 | 2017 ± 380 |
| 19,20-DiHDPA | 1014 ± 143 | 763 ± 117 | 512 ± 179 | 483 ± 29 |

**Supplementary Table 3.** Lipid mediators in 48h blisters. Concentrations in pg/ml (mean ± sem).

|  | HC | UC | UC5ASA | HC 5ASA |
| --- | --- | --- | --- | --- |
| **Eicosanoids** |  |  |  |  |
| PGD2 | 293 ± 54 | 579 ± 179 | 372 ± 146 | 227 ± 48 |
| PGE1 | 150 ± 24 | 145 ± 38 | 117 ± 53 | 276 ± 84 |
| 13,14-dihydro-15-keto-PGE1 | 6 ± 4 | 7 ± 7 | 6 ± 6 | 23 ± 23 |
| PGE2 | 1168 ± 164 | 1426 ± 256 | 1161 ± 272 | 1570 ± 348 |
| 13,14-dihydro-15-keto-PGE2 | 148 ± 27 | 251 ± 49 | 216 ± 49 | 150 ± 26 |
| D12 PGJ2 | 704 ± 121 | 670 ± 111 | 727 ± 124 | 584 ± 92 |
| 15-deoxy-D12,14-PGJ2 | 171 ± 58 | 103 ± 24 | 118 ± 26 | 83 ± 12 |
| PGF1α | 135 ± 29 | 104 ± 24 | 83 ± 31 | 75 ± 56 |
| 6-keto-PGF1α | 138 ± 27 | 94 ± 16 | 66 ± 22 | 89 ± 30 |
| PGF2α | 1331 ± 215 | 1214 ± 204 | 1184 ± 391 | 521 ± 76 |
| 13,14-dihydro-15-keto-PGF2α | 0 | 0 | 0 | 0 |
| TXB2 | 742 ± 190 | 669 ± 197 | 271 ± 107 | 395 ± 66 |
| **Hydroxy lipids** |  |  |  |  |
| 4-HDHA | 251 ± 52 | 245 ± 123 | 328 ± 113 | 515 ± 89 |
| 13-HDHA | 0 | 18 ± 18 | 0 | 0 |
| 14-HDHA | 111 ± 34 | 53 ± 38 | 14 ± 14 | 87 ± 53 |
| 17-HDHA | 79 ± 79 | 563 ± 371 | 0 | 0 |
| 20-HDHA | 73 ± 36 | 58 ± 38 | 44 ± 44 | 0 |
| 11,12-DHET | 159 ± 17 | 129 ± 28 | 167 ± 24 | 209 ± 35 |
| 14,15-DHET | 231 ± 10 | 225 ± 40 | 244 ± 43 | 211 ± 61 |
| 5-HETE | 23 ± 15 | 42 ± 27 | 42 ± 28 | 139 ± 63 |
| 11-HETE | 225 ± 22 | 165 ± 30 | 176 ± 35 | 129 ± 26 |
| 12-HETE | 4970 ± 1895 | 2127 ± 444 | 2038 ± 354 | 3465 ± 1349 |
| 15-HETE | 815 ± 74 | 933 ± 445 | 497 ± 173 | 670 ± 130 |
| 20-HETE | 161 ± 84 | 234 ± 149 | 0 | 169 ± 169 |
| 15-HETrE | 468 ± 68 | 594 ± 191 | 335 ± 74 | 529 ± 149 |
| 11(12)-EET | 103 ± 50 | 21 ± 21 | 46 ± 30 | 50 ± 33 |
| 12-HEPE | 169 ± 92 | 13 ± 13 | 5 ± 5 | 40 ± 24 |
| 18-HEPE | 38 ± 25 | 0 | 0 | 0 |
| 9-HODE | 3950 ± 513 | 4501 ± 517 | 4205 ± 739 | 2653 ± 612 |
| 13-HODE | 9210 ± 1122 | 10221 ± 1613 | 9679 ± 1542 | 6491 ± 1310 |
| 9-OxoODE | 3362 ± 404 | 4387 ± 238 | 6023 ± 978 | 2005 ± 170 |
| 13-OxoODE | 1862 ± 319 | 2899 ± 406 | 3955 ± 507 | 1442 ± 200 |
| 9-HOTrE | 532 ± 121 | 625 ± 182 | 482 ± 200 | 1453 ± 371 |
| 13-HOTrE | 288 ± 34 | 192 ± 60 | 158 ± 68 | 387 ± 86 |
| 9(10)-EpOME | 1386 ± 378 | 1069 ± 256 | 1462 ± 337 | 1632 ± 219 |
| 12(13)-EpOME | 1719 ± 452 | 1451 ± 319 | 1918 ± 220 | 1562 ± 173 |
| Trans-EKODE | 964 ± 142 | 1195 ± 145 | 1340 ± 332 | 723 ± 218 |
| 9,10-DiHOME | 1231 ± 232 | 1326 ± 206 | 1566 ± 383 | 1728 ± 272 |
| 12,13-DiHOME | 1630 ± 317 | 1444 ± 395 | 1979 ± 574 | 2078 ± 480 |
| 19,20-DiHDPA | 694 ± 85 | 733 ± 129 | 759 ± 208 | 347 ± 63 |

